## Supplementary Information for "Heritable epigenetic changes are constrained by the dynamics of regulatory architectures"

Antony M Jose[1]*

**Affiliations:**

[1]University of Maryland, College Park, MD, USA.

**Content:**

Supplementary methods
Supplementary Figures and Figure Legends (Fig. S1 – S14)
Supplementary Movie Legends (Movie. S1 – S13)

### SUPPLEMENTARY METHODS

**Analysis of simple heritable regulatory architectures.** The dynamics of the 26 heritable regulatory architecture (HRAs in Fig. 1) can be described by systems of ordinary differential equations. The rate of change of each entity ($x$, $y$, or $z$ for each HRA in Fig. 1) can be written by aggregating positive and negative contributions from the other sensors (e.g., $k_{xy}.y - k_{xz}.z$ if $x$ is positively regulated by $y$ and negatively regulated by $z$) with loss of each entity described with a turnover term (e.g., $-T_x.x$ for $x$). To ensure the concentrations of all entities remain non-negative, as expected in real living systems, the equations are bounded to be applicable only when the values of the changing entity is greater than zero. These equations can then be used to derive other equations and inequalities of interest.

<u>*Steady-state relationships*</u>. At steady state, each architecture results in relative amounts of each entity ($x_0, y_0, z_0$) that could define a 'phenotype'. These relative concentrations can be derived by setting the rate of change of all entities to zero.

For the heritable regulatory architecture A,

$$\frac{dx}{dt} = k_{xy}.y - T_x.x, \qquad \forall\ (x + dx) > 0, \text{else } (x + dx) = 0$$

$$\frac{dy}{dt} = k_{yx}.x - T_y.y, \qquad \forall\ (y + dy) > 0, \text{else } (y + dy) = 0$$

where, $k_{xy}$ is the rate constant for the production of $x$ promoted by $y$; $k_{yx}$ is the rate constant for the production of $y$ promoted by $x$; $T_x$ is the rate of turnover of $x$; and $T_y$ is the rate of turnover of $y$.

i.e., $\dot{X} = A.X$, where $\dot{X} = \begin{bmatrix} \dot{x} \\ \dot{y} \end{bmatrix}$; A = $\begin{bmatrix} -T_x & k_{xy} \\ k_{yx} & -T_y \end{bmatrix}$; $X = \begin{bmatrix} x \\ y \end{bmatrix}$

At steady state,

$$-T_x.x + k_{xy}.y = 0$$

$$k_{yx}.x - T_y.y = 0$$

i.e., $A.X = 0$, where A = $\begin{bmatrix} -T_x & k_{xy} \\ k_{yx} & -T_y \end{bmatrix}$; $X = \begin{bmatrix} x_0 \\ y_0 \end{bmatrix}$, which has solutions that satisfy:

$$\frac{x_0}{y_0} = \frac{k_{xy}}{T_x} = \frac{T_y}{k_{yx}}$$

Equations for steady state and the resulting solutions for the other HRAs can be similarly derived (see previous version of the paper for details).

<u>*Steady states with loss of all entities to complex formation at a constant rate*</u>. A common way in which entities change in living systems is through the formation of intermolecular complexes that then interact with different entities to perform different functions. If the same number of molecules per unit time (γ) are lost for all entities (e.g., through incorporation into a 1:1:1 stoichiometric complex), then each entity needs to grow at the same rate (γ) to maintain steady state ($x_0, y_0, z_0$).

For the heritable regulatory architecture A,

$$k_{xy}.y - T_x.x = \gamma$$

$$-T_y.y + k_{yx}.x = \gamma$$

i.e., $A.X = B$, where A = $\begin{bmatrix} -T_x & k_{xy} \\ k_{yx} & -T_y \end{bmatrix}$; $X = \begin{bmatrix} x_0 \\ y_0 \end{bmatrix}$; $B = \begin{bmatrix} \gamma \\ \gamma \end{bmatrix}$

The solution is given by $X = A^{-1}.B$

(For A = $\begin{bmatrix} a & b \\ c & d \end{bmatrix}$, $A^{-1} = \frac{1}{ad-bc}\begin{bmatrix} d & -b \\ -c & a \end{bmatrix}$)

$$X = \frac{1}{T_x.T_y - k_{xy}.k_{yx}}\begin{bmatrix} -T_y & -k_{xy} \\ -k_{yx} & -T_x \end{bmatrix}.\begin{bmatrix} \gamma \\ \gamma \end{bmatrix}$$

$$i.e., \begin{bmatrix} x_0 \\ y_0 \end{bmatrix} = \frac{\gamma}{k_{xy}.k_{yx} - T_x.T_y}\begin{bmatrix} T_y + k_{xy} \\ T_x + k_{yx} \end{bmatrix}$$

To similarly identify the rates of growth for the other heritable regulatory architectures (B to Z) under a constant rate of loss for all entities, inverses for the 3x3 matrices can be used (see previous version of the paper for details). If all molecules are diluted through cell division (typically one cell dividing to give two), then for maintaining steady state on average, each molecule needs to accumulate to 2x the average steady-state value per cell cycle (See Fig. 4 for simulations that include cell divisions).

*Response to genetic loss of an entity*. Loss of an entity (typically the RNA or protein product of a gene) through a genetic mutation is a common perturbation used for analyzing living systems.

For the heritable regulatory architecture A,

When $x$ is lost,

$$\dot{y} = -T_y.y$$

Which has the solution,

$y = y_0.e^{-T_y t}$, i.e., the concentration of $y$ undergoes exponential decay through turnover from its steady-state value ($y_0$).

Similarly, when $y$ is lost,

$$x = x_0.e^{-T_x t}$$

For all other architectures, loss of one entity can result in different dynamics of the other two entities ($\alpha$ and $\beta$, say) depending on their regulatory interactions. The equations for their dynamics is given by a pair of differential equations that can be coupled.

i.e., $\dot{X} = A.X$, where $\dot{X}$ = $\begin{bmatrix} \dot{\alpha} \\ \dot{\beta} \end{bmatrix}$; A = $\begin{bmatrix} a & b \\ c & d \end{bmatrix}$; $X = \begin{bmatrix} \alpha \\ \beta \end{bmatrix}$.

The exact equations that result upon loss of each entity in each regulatory architecture can be derived for each HRA (see previous version of the paper for details) and potentially be used to distinguish the different architectures through genetic experiments (e.g., knockout of individual genes using genome editing). Since for the steady state of each architecture, there are a maximum of three equations, a maximum of three variables among rates of production ($k_{xy}, k_{yx}$ etc.), rates of turnover ($T_x, T_y, T_z$), and the steady-state concentrations ($x_0, y_0, z_0$) are constrained. Changes in all entities after each loss can be determined for all architectures through simulations by choosing random values for the unconstrained parameters (Figures S2 to S8, left).

*Response to epigenetic change*. Analytic expressions for heritable epigenetic change after reducing the levels of a sensor from steady state are derived below for the simplest of heritable regulatory architectures 'A' (Fig. 1).

The dynamics of two entities ($x$ and $y$) that mutually promote each other's production is given by a pair of differential equations that are coupled.

i.e., $\dot{X} = A.X$, where $\dot{X} = \begin{bmatrix}\dot{x} \\ \dot{y}\end{bmatrix}$; A = $\begin{bmatrix}a & b \\ c & d\end{bmatrix}$; $X = \begin{bmatrix}x \\ y\end{bmatrix}$ and $a = -T_x, b = k_{xy}, c = k_{yx}, \; d = \; -T_y$

The general solution for the concentrations $x(t)$ and $y(t)$ are:

$$x(t) = \frac{K_1}{2m}\left(e^{0.5t(a+d-m)}(e^{tm}(a-d+m)+m+d-a)\right) - \frac{bK_2}{m}(e^{0.5t(a+d-m)} - e^{0.5t(a+d+m)})$$

$$y(t) = \frac{K_2}{2m}\left(e^{0.5t(a+d-m)}(e^{tm}(d-a+m)+m-d+a)\right) - \frac{cK_1}{m}(e^{0.5t(a+d-m)} - e^{0.5t(a+d+m)})$$

where, $m = \sqrt{a^2 - 2ad + 4bc + d^2}$ and, $K_1$and $K_2$are constants.

Since the system was already at steady state before the perturbation, $T_xT_y = k_{xy}k_{yx}$,

$$m = \sqrt{{T_x}^2 - 2T_xT_y + 4k_{xy}k_{yx} + {T_y}^2} = T_x + \; T_y$$

Substituting the value of $m$ in the equations above and simplifying yields,

$x(t) = \frac{K_1.T_y + K_2.k_{xy}}{T_x + T_y} + \frac{K_1.T_x - K_2.k_{xy}}{T_x + T_y}.e^{-(T_x+T_y)t}$ and $y(t) = \frac{K_2.T_x + K_1.k_{yx}}{T_x + T_y} + \frac{K_2.T_y - K_1.k_{yx}}{T_x + T_y}.e^{-(T_x+T_y)t}$

Let $d_x$ be the reduction in $x$ (reduction-of-function) needed to observe a defect when $x_0$ is the steady-state value before perturbation. That is, $d_x.x_0$ is not sufficient for the function of $x$ in a living system, where $d_x < 1$. Let $x$ be perturbed to $x_p = p.d_x.x_0 \; \neq 0$ from $t = 0$ until $t = t_p$, where $p < 1$. For heritable epigenetic changes using reduction-of-function perturbations ($d_x < 1$ and/or $d_y < 1$), which preserve the architecture at a new steady state: $x_{ps} < d_x.x_0$ and $y_{ps} < d_y.y_0$.

To determine the concentration of $y$ at the end of the perturbation ($y_p$) the equation $\dot{y} = \; x_p.k_{yx} - T_y.y$ can be solved using $y(t) \; = \; y_0$ at $t = \; 0$. The general solution of the equation is given by,

$$y(t) = \; \frac{x_p.k_{yx}}{T_y} + C_1.e^{-T_yt}$$

Substituting for $y(0) = y_0$ at $t = 0$, and rearranging gives $C_1 = \; \frac{y_0.T_y \; - x_p.k_{yx}}{T_y}$. Thus, at the end of the perturbation (i.e., at $t_p$),

$$y(t_p) = y_p = \frac{x_p.k_{yx}}{T_y} + \; (\frac{y_0.T_y \; - x_p.k_{yx}}{T_y}).e^{-T_yt_p}$$

The new steady states ($x_{ps}$ and $y_{ps}$) will be reached from the initial concentrations of $x_p$ and $y_p$.

Therefore, to determine the new steady state, the initial values of $x_p$ and $y_p$ can be used at new $t = 0$ to get the values for the constants $K_1$ and $K_2$.

$$x_p = \frac{K_1.T_y + K_2.k_{xy}}{T_x + T_y} + \frac{K_1.T_x - K_2.k_{xy}}{T_x + T_y}.e^{-(T_x+T_y)t=0}$$

$$y_p = \frac{K_2.T_x + K_1.k_{yx}}{T_x + T_y} + \frac{K_2.T_y - K_1.k_{yx}}{T_x + T_y}.e^{-(T_x+T_y)t=0}$$

Which simplifies to,

$$x_p = \frac{K_1.T_y + K_2.k_{xy}}{T_x + T_y} + \frac{K_1.T_x - K_2.k_{xy}}{T_x + T_y}$$

$$y_p = \frac{K_2.T_x + K_1.k_{yx}}{T_x + T_y} + \frac{K_2.T_y - K_1.k_{yx}}{T_x + T_y}$$

Solving for each,

$$K_1 = x_p$$

$$K_2 = y_p$$

To obtain the new steady state value $x_{ps}$, set $t = \infty$ in the equation using the above constants.

$$x_{ps} = \frac{x_p.T_y + y_p.k_{xy}}{T_x + T_y}$$

$$y_{ps} = \frac{y_p.T_x + x_p.k_{yx}}{T_x + T_y}$$

Consider the equality that is the threshold for observing heritable epigenetic effects,

$$x_{ps} = \frac{x_p.T_y + y_p.k_{xy}}{T_x + T_y} = d_x.x_0$$

Substituting for $x_p$ and simplifying yields,

$$y_p.k_{xy} = d_x.x_0.T_x + d_x.x_0.T_y(1-p)$$

Substituting for $y_p$

$$(\frac{x_p.k_{yx}}{T_y} + (\frac{y_0.T_y - x_p.k_{yx}}{T_y}).e^{-T_y t_p}).k_{xy} = d_x.x_0.T_x + d_x.x_0.T_y(1-p)$$

Collecting exponential terms,

$$e^{-T_y t_p}(\frac{y_0.k_{xy}.T_y - p.d_x.x_0.k_{yx}.k_{xy}}{T_y}) = d_x.x_0.T_x + d_x.x_0.T_y(1-p) - \frac{p.d_x.x_0.k_{yx}.k_{xy}}{T_y}$$

$$e^{-T_y t_p} = \frac{d_x.x_0.T_x.T_y + d_x.x_0.T_y.T_y.(1-p) - p.d_x.x_0.k_{yx}.k_{xy}}{y_0.k_{xy}.T_y.T_y - p.d_x.x_0.k_{yx}.k_{xy}}$$

$$e^{-T_y t_p} = \frac{d_x.x_0.T_x.T_y + d_x.x_0.T_y.T_y - p.d_x.x_0.(k_{yx}.k_{xy} + T_y.T_y)}{y_0.k_{xy}.T_y - p.d_x.x_0.k_{yx}.k_{xy}}$$

Dividing numerator and denominator with $T_y$,

$$e^{-T_y t_p} = \frac{d_x.x_0.T_x + d_x.x_0.T_y - p.d_x.x_0.(\frac{k_{yx}.k_{yx}}{T_y} + T_y)}{y_0.k_{xy} - p.d_x.x_0.\frac{k_{yx}.k_{yx}}{T_y}}$$

At steady state, the ratio $\frac{x(t)}{y(t)}$ will be independent of the concentrations of $x$ and $y$. That is, $\frac{x_{ps}}{y_{ps}} = \frac{x_0}{y_0} = \frac{k_{xy}}{T_x} = \frac{T_y}{k_{yx}}$. Therefore, these equalities can be used to simplify the above equations. Substituting $\frac{k_{yx}.k_{yx}}{T_y} = T_x$,

$$e^{-T_y t_p} = \frac{d_x.x_0.T_x + d_x.x_0.T_y - p.d_x.x_0.(T_x + T_y)}{y_0.k_{xy} - p.d_x.x_0.T_x}$$

Substituting $y_0.k_{xy} = x_0.T_x$,

$$e^{-T_y t_p} = \frac{d_x.x_0.T_x + d_x.x_0.T_y - p.d_x.x_0.(T_x + T_y)}{x_0.T_x - p.d_x.x_0.T_x}$$

Dividing numerator and denominator by $x_0.T_x$

$$e^{-T_y t_p} = \frac{d_x + d_x.\frac{T_y}{T_x} - p.d_x.(1 + \frac{T_y}{T_x})}{1 - p.d_x}$$

Simplifying,

$$e^{-T_y t_p} = \frac{(1 + \frac{T_y}{T_x}).(1 - p).d_x}{1 - p.d_x}$$

Dividing numerator and denominator by $d_x$

$$e^{-T_y t_p} = \frac{(1 + \frac{T_y}{T_x}).(1 - p)}{\left(\frac{1}{d_x} - p\right)}$$

Taking the $log_e$ on both sides,

$$-T_y t_p = ln\left[\frac{\left(1 + \frac{T_y}{T_x}\right).(1 - p)}{\frac{1}{d_x} - p}\right]$$

i.e.,

$$t_p = \frac{1}{T_y} ln \left[ \frac{\frac{1}{d_x} - p}{\left(1 + \frac{T_y}{T_x}\right).(1-p)} \right]$$

Dividing numerator and denominator within the antilogarithm by $p$,

$$t_p = \frac{1}{T_y} ln \left[ \frac{\frac{1}{d_x.p} - 1}{\left(\frac{1}{p} - 1\right)\left(1 + \frac{T_y}{T_x}\right)} \right]$$

This equation relates the duration of a perturbation ($t_p$) and the extent of the perturbation ($p < 1$ for loss-of-function) beyond the threshold that causes a defect in the function of $x$ (i.e., $d_x$). Increasing the duration of the perturbation beyond $t_p$ for a given extent of perturbation ($p$) will result in heritable epigenetic change where the steady-state levels of both interactors are insufficient for appropriate function.

Similarly, the minimal duration of perturbation for heritable epigenetic changes through a defect in the function of $y$ is given by,

$$t_p > \frac{1}{T_x} ln \left[ \frac{\frac{1}{d_y.p} - 1}{\left(\frac{1}{p} - 1\right)\left(1 + \frac{T_x}{T_y}\right)} \right]$$

These inequalities were verified using numerical simulations (see 'HRA_A_ tp_analytical_expression_check.py') and additional HRAs were similarly simulated to gain intuitions about the consequences of epigenetic reduction in the levels of entities (Figures S3 to S9).

**Simulation of simple ESP systems.** A model was created in NetLogo to simulate entity-sensor-property systems and their evolution across generations for exploring regulatory architectures (ESP_systems _explorer_v1.nlogo). A variety of regulatory architectures were simulated using this model to identify ones with some architecture that persisted for 250 generations with or without perturbations (e.g., 78285 out of 225000 tested using the experiment "ESP_origins_2-16_mols" described under behaviorspace). Each stable ESP system can be further analyzed in detail using the related ESP_systems_single_system_explorer_v1. All randomly chosen values for parameters in the regulatory architectures that lead to stability, i.e., heritable for many generations, can be recreated because the random-seed is set for each run using the behaviorspace-run-number. The behaviorspace-run-number serves as the 'system-id' in the related ESP_systems_single_system_explorer_v1.nlogo.

For each system, entities/sensors were defined as 'turtles' with 3 variables that stored its attributes:

(1) val – a variable for storing an entity/sensor's current 'property value' (e.g., concentration, % conformational change, sequence, etc.). It changes throughout the simulation and is always represented in architectures as the size of the circle for each entity after scaling it relative to all extant entities.

(2) property – a variable for storing the steps of change by which values (i.e., the 'val' above) can change if a positive or negative interaction crosses the threshold for change. For these simulations, it is characteristic of the entities/sensors themselves and does not change as the system evolves through interactions, which introduces the simplification that every sensor sees the same property of a given entity/sensor.
(3) inactive-fraction – a variable for storing the fraction of entity/sensor not available for regulatory interactions at each time step (tick) because of processes like protein folding, compartmentalization, diffusion, etc. This is a characteristic of each entity/sensor that is randomly chosen at the beginning of the simulation and does not change during the simulation.

The regulatory interactions in each system were specified using 'links' that were weighted to indicate the threshold required for the regulatory interaction and colored to indicate the nature of the regulation. Specifically, the links have two variables:
(1) weight - a variable that indicates the number of sensors needed to change one unit of property for each entity. This parameter is characteristic of each regulatory interaction and captures the threshold needed for transmission of change. For display, the thickness of the regulatory link is set to be 0.5 - weight / 20. Thus, a lower threshold for transmission is represented as a wider link.
(2) color - a variable that indicates whether the regulatory interaction is positive (grey) or negative (black).

Parameters that were varied in the exploration of ESP systems were:
(1) molecule-kinds, which was the number of entities/sensors that are part of the regulatory architecture.
(2) perturb-kind (none, lof, or gof), which was a chooser for perturbing a random entity/sensor every ~50 generations for 2.5 generations by increasing (gof) or decreasing (lof) its value (i.e., concentration/number) by two fold of the maximal or minimal values, respectively, of all the entities/sensors.
(3) perturb-phase, which was the precise timing for starting the periodic perturbations (e.g., 0 = starting @ tick 100; 1 = starting @ tick 101; 2 = starting @ tick 102; 3 = starting @ tick 103; 4 = starting @ tick 104)

Additional parameters, which were not varied in the exploration of ESP systems were:
(1) cycle-time, which was set at 2 and represented the timing in ticks for each generation.
(2) link-chance, which was set at 50% and gave the probability that any two entities/sensors will interact when the system is set up at the beginning of the simulation.
(3) positive-interactions, which was set at 50% and gave the probability that a regulatory interaction is positive.
(4) max-molecules, which was set at 500 and was the maximal number of total molecules at the start of the simulation.
(5) stasis-level, which was set at 5000 and was the number of molecules that arrests growth until molecules get diluted upon cell division.
(6) max-ever-molecules, which was set at 500000 was the maximal number of molecules of all kinds put together that can be within any system at any time. This limit simulates living systems existing in a finite environment.

Monitors reporting behaviorspace-run-number (system-id), generation number, total molecules, the number of generations of stability for considering a regulatory architecture stable (stability gen), stable generations since last instability, the value of the perturbed entity (perturb-value), the phase of the perturbation (perturb-phase), the duration of each perturbation (perturbtime), the frequency of the perturbations (perturb-freq) and the identity of the perturbed entity (perturbed node) were included in the interface.

For simulating changes over time, this model used a combination of deterministic and stochastic functions. The values of each entity/sensor changes at each tick using a deterministic equation: val @ t+1 = val @ t + sum of inputs from all sensors. The change in value contributed by each sensor for a given entity = round((property of entity) x (value of sensor) x (1 - inactive-fraction of sensor) / (weight of regulatory link)). In other words, change = round(*k* x (value of sensor)), where *k* is a different constant for each sensor of each entity and round indicates rounding to the nearest integer. For positive regulators (link color grey), this change in value was added and for negative regulators (link color black), it was subtracted. The order of operation on the entities/sensors varies with every tick. After every two ticks, only about half the number of each entity/sensor was kept using a random number generator to simulate dilution and random partitioning upon cell division.

The value of each entity/sensor was plotted relative to the most abundant entity/sensor. This profile at each time point can be considered as the 'phenotype' of the system. These scaled values are also used to depict each entity/sensor in the regulatory architecture at each time point.

**Exploration of simple ESP systems.** To gain intuitions by exploring and perturbing simulated ESP systems, several interactive features were added to the ESP simulator (Fig. S11, Movie S1). These include parameters that control the setup and running of randomly generated ESP systems by specifying the probability of regulatory interactions in the system (link-chance, Fig. S11A), the probability of positive versus negative interactions (positive-interactions, Fig. S11A), the maximum number of molecules at the start of the simulation (max-molecules, Fig. S11A), the maximum number of molecules that will arrest growth until dilution through cell divisions to simulate depletion of raw materials or energy (stasis-level, Fig. S11A), maximum number of molecules in total reflecting the limited space occupied by living systems (max-ever-molecules, Fig. S11A), and duration of the cell cycle (cycle-time, Fig. S11A). Particular systems can be re-established and re-simulated by setting the random number seed that is used for controlling all stochastic steps (system-id, Fig. S11B) and by additionally specifying the above parameters along with the number of entities/sensors at the start of the simulation (molecule-kinds, Fig. S11B). For each such system, the number of entities that can increase or decrease at one time was set to be characteristic of each entity/sensor (unit change in property value, i.e., number) and the number of sensors needed to change one unit of each entity/sensor was set to be characteristic of each regulatory interaction (thickness of link increases with increasing sensitivity of regulation). Periodic loss-of-function or gain-of-function perturbations (perturb-kind, Fig. S11B) can be set up to begin in five different phases relative to the start of the simulation (perturb-phase [0, 1, 2, 3 or 4], Fig. S11B). Perturbations that can be made during the simulation include changing the number of molecules of any entity/sensor (change-a-node, Fig. S11C), adding an entity/sensor (add-a-node, Fig. S11C), removing an entity/sensor (remove-a-node, Fig. S11C), removing a particular regulatory interaction (remove-link-x-y, Fig. S11C), removing a random regulatory interaction (remove-a-link, Fig. S11C), and changing the strength of a regulatory input (link-hold, Fig. S11C). Finally, a reporter for any entity/sensor (add-a-reporter, Fig. S11D) can be set up that either perfectly or partially interacts with all its regulators (perfect?, Fig. S11D). A perfect reporter of an entity/sensor receives the same regulatory input as the entity/sensor of interest does. An imperfect reporter of an entity receives input from the same sensors as the entity/sensor of interest, but the polarity and strength of the input can vary. Regulatory outputs of the entity/sensor are not recreated for any reporter. As these changes are being made, both the regulatory architecture (Fig. S11E, *top*), which is re-drawn if the levels of any entity/sensor reaches zero, and the ‘phenotype’ as captured by the profile of relative concentrations of entities/sensors (Fig. S11E, *bottom*) can be observed.

### SUPPLEMENTARY FIGURES AND FIGURE LEGENDS

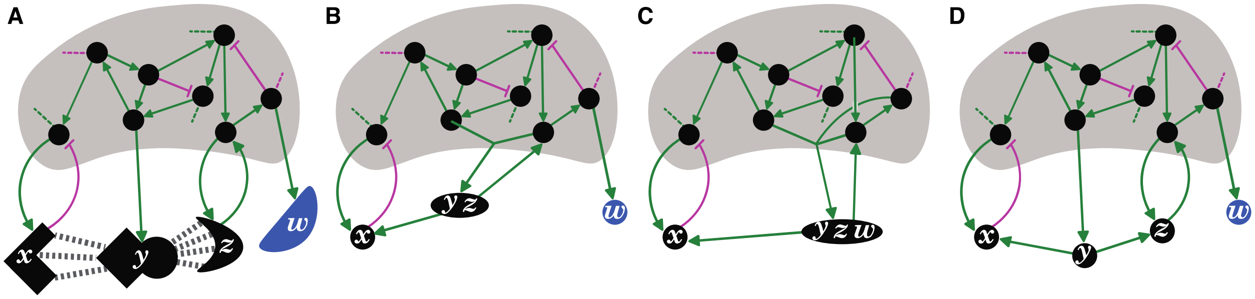

**Figure S1. Entity-Sensor-Property systems provide a principled way of parsing regulators and their interactions in living systems.** (A) Schematic of regulatory interactions in a living system, highlighting incomplete knowledge, but including some regulators that detect the shape(s) of others. Entities that act as sensors (black circles and black shapes) by providing regulatory input in response to changes in other entities or that do not provide any regulatory input (blue shape), interactions that promote (green arrows) or inhibit (magenta bar) a property of downstream entities/sensors, interactions with unknown entities/sensors (dotted lines), and the unknown larger network (grey shading) are depicted. (B and C) Two ways of parsing the interactors that combine some regulators together (*y* and *z* in (B), and *y*, *z*, and *w* in (C)) and therefore do not reflect the natural properties salient to the system in (A). (D) Deduced regulatory architecture with sensors (*x*, *y*, *z*; red) and entities (*w*; blue) parsed to better reflect the system depicted in (A). Progression from the depiction in (B) or (C) to that in (D) requires experiments that consider the separable entities (*x*, *y*, *z* and *w*), sensors (*x*, *y* and *z*), and the sensed properties (*y*'s square edges for sensor *x*, and its curved surfaces for sensor *z*) that are relevant for the system.

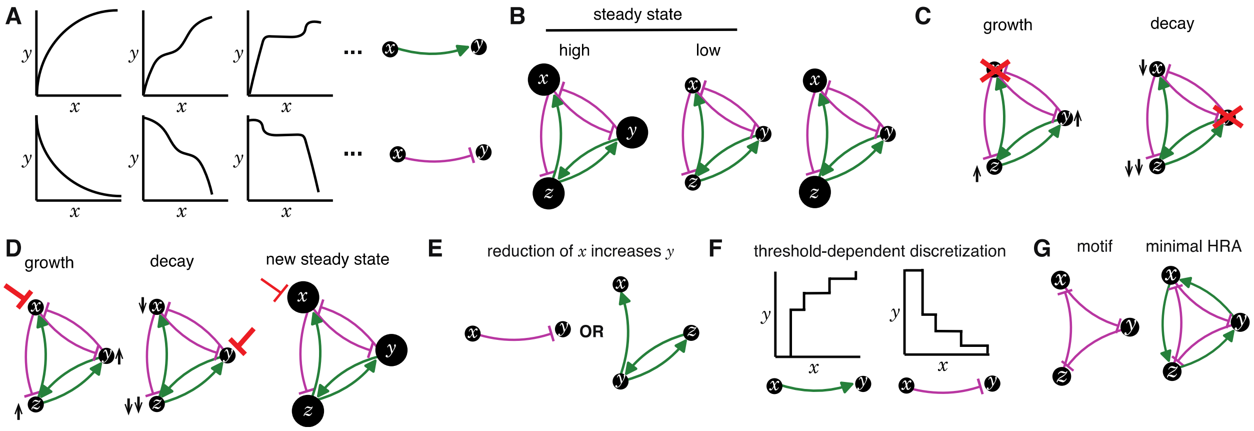


**Figure S2. Illustrations of key concepts.** (A) Regulatory interactions can lead to a variety of dependencies between two entities. Alternative relationships (*left*) when *x* promotes *y* (*top*) or when *x* inhibits *y* (*bottom*) and their generic representations (*right*; green arrow = promote and magenta bar = inhibit). In general, the relationship between any two entities in a living system requires empirical investigation. (B) The same regulatory architecture (HRA 'Y' from Fig. 1) can be present at different states with relatively high (*left*) or proportionally low (*middle*) levels of all entities (areas of circles) at steady state, or with unregulated levels of different entities (*right*) away from steady state (e.g., soon after a perturbation). (C) After permanent loss of an entity (red x), the remaining entities can show uncontrolled growth (*left*, up arrow) or eventual decay (*right*, down arrow) depending on the residual architecture. (D) After transient reduction in the levels of an entity (red bar), the remaining entities can show uncontrolled growth (*left*), eventual decay (*middle*), or recovery to a new steady state level (*right*) depending on the residual architecture and the strength/duration of the perturbation. (E) Two equivalent inferences based on an observation after a perturbation whereby reduction in *x* increases *y*. Either *x* inhibits *y* (*left*) or *y* promotes *x* and itself via *z* (*right*). (F) The thresholds required for interactions between entities discretize the changes caused by promotions (*left*) or inhibitions (*right*) in living systems. (G) An regulatory motif composed of three repressors (*left*) needs at least three positive regulatory interactions to be heritable (*right*).

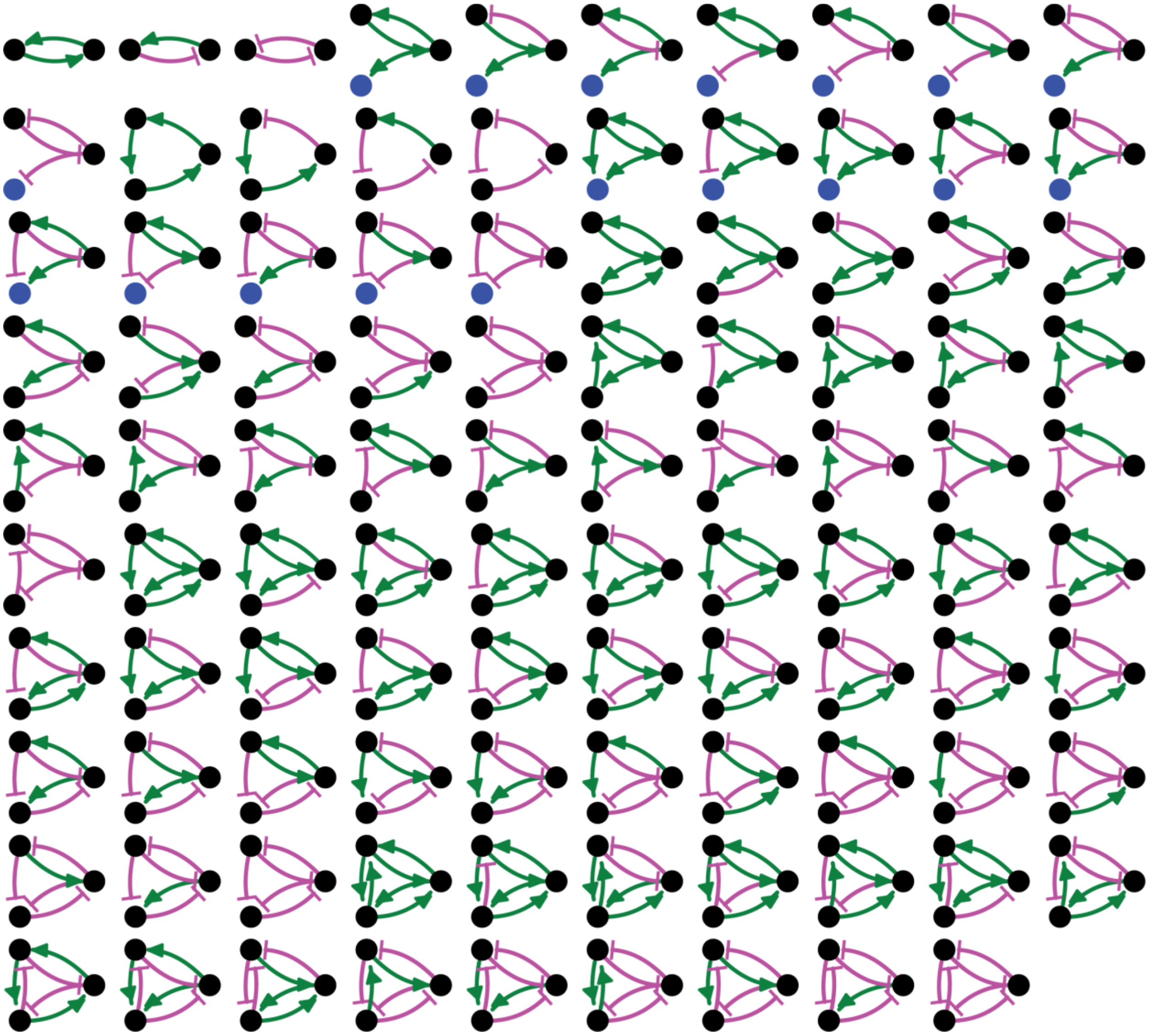

**Figure S3. Adding regulation to the 8 simple heritable architectures generates 99 regulated architectures, not all of which are heritable.** Entities that act as sensors (black circles) or that do not provide any regulatory input (blue circles), positive (green arrows) and negative (magenta bar) regulatory interactions are indicated.

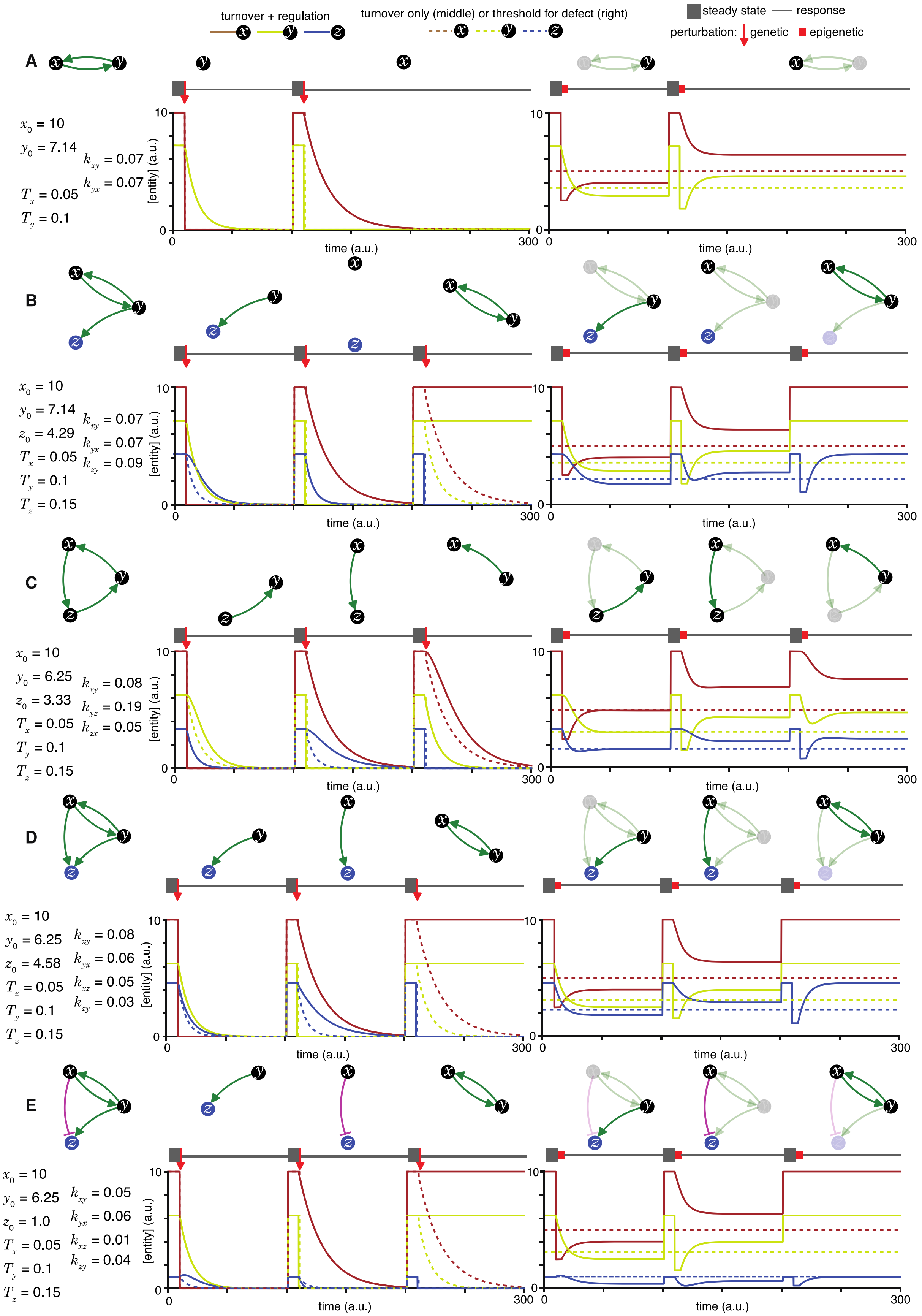
turnover + regulation
turnover only (middle) or threshold for defect (right)
steady state
response
perturbation: genetic
epigenetic
A
$x_0$ = 10
$y_0$ = 7.14
$k_{xy}$ = 0.07
$k_{yx}$ = 0.07
$T_x$ = 0.05
$T_y$ = 0.1
B
$x_0$ = 10
$y_0$ = 7.14
$z_0$ = 4.29
$k_{xy}$ = 0.07
$k_{yx}$ = 0.07
$k_{zy}$ = 0.09
$T_x$ = 0.05
$T_y$ = 0.1
$T_z$ = 0.15
C
$x_0$ = 10
$y_0$ = 6.25
$z_0$ = 3.33
$k_{xy}$ = 0.08
$k_{yz}$ = 0.19
$k_{zx}$ = 0.05
$T_x$ = 0.05
$T_y$ = 0.1
$T_z$ = 0.15
D
$x_0$ = 10
$y_0$ = 6.25
$z_0$ = 4.58
$k_{xy}$ = 0.08
$k_{yx}$ = 0.06
$k_{xz}$ = 0.05
$k_{zy}$ = 0.03
$T_x$ = 0.05
$T_y$ = 0.1
$T_z$ = 0.15
E
$x_0$ = 10
$y_0$ = 6.25
$z_0$ = 1.0
$k_{xy}$ = 0.05
$k_{yx}$ = 0.06
$k_{xz}$ = 0.01
$k_{zy}$ = 0.04
$T_x$ = 0.05
$T_y$ = 0.1
$T_z$ = 0.15
[entity] (a.u.)
time (a.u.)
0
10
300

**Figure S4. Heritable regulatory architectures with one loop.** Loss-of-function perturbations of each entity (*x*, *y*, or *z*, if present) in architectures (*left top*) characterized by sets of parameters that support a steady state (*left bottom*) are illustrated. The behaviour of residual architectures after permanent or genetic (*middle*) and transient or epigenetic (*right*) changes are illustrated. Period of steady state (thick grey line), the point of genetic change (red arrow), duration of epigenetic reduction (red bar, for a duration $t_p = 5$ (a.u); with the threshold for observing a defect $d = 0.5$; and an extent of perturbation beyond the threshold $p = 0.5$), and duration of recovery after perturbation (thin grey line) were simulated. Architectures are depicted as in Figure 1 (A, B, C, D, and E depict the heritable regulatory architectures A, B, C, D, and E, respectively) with transient reductions in an entity and associated interactions depicted using lighter shades. Dotted lines indicate unregulated turnover (in *middle*) or thresholds for observing defects upon reduction in levels of an entity (in *right*).

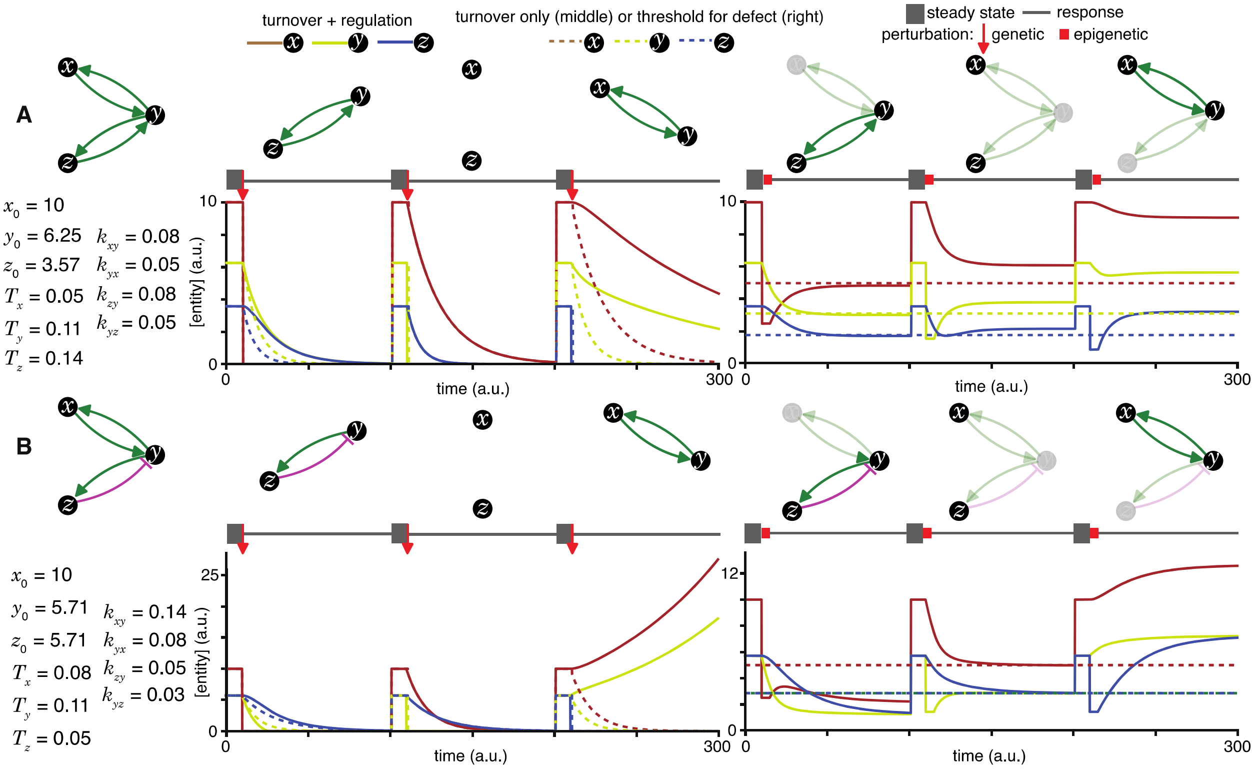


**Figure S5. Heritable regulatory architectures with two loops and a shared node.** Architectures and their responses to perturbations are depicted as in Figure S4 (A and B depict the heritable regulatory architectures F and G, respectively).

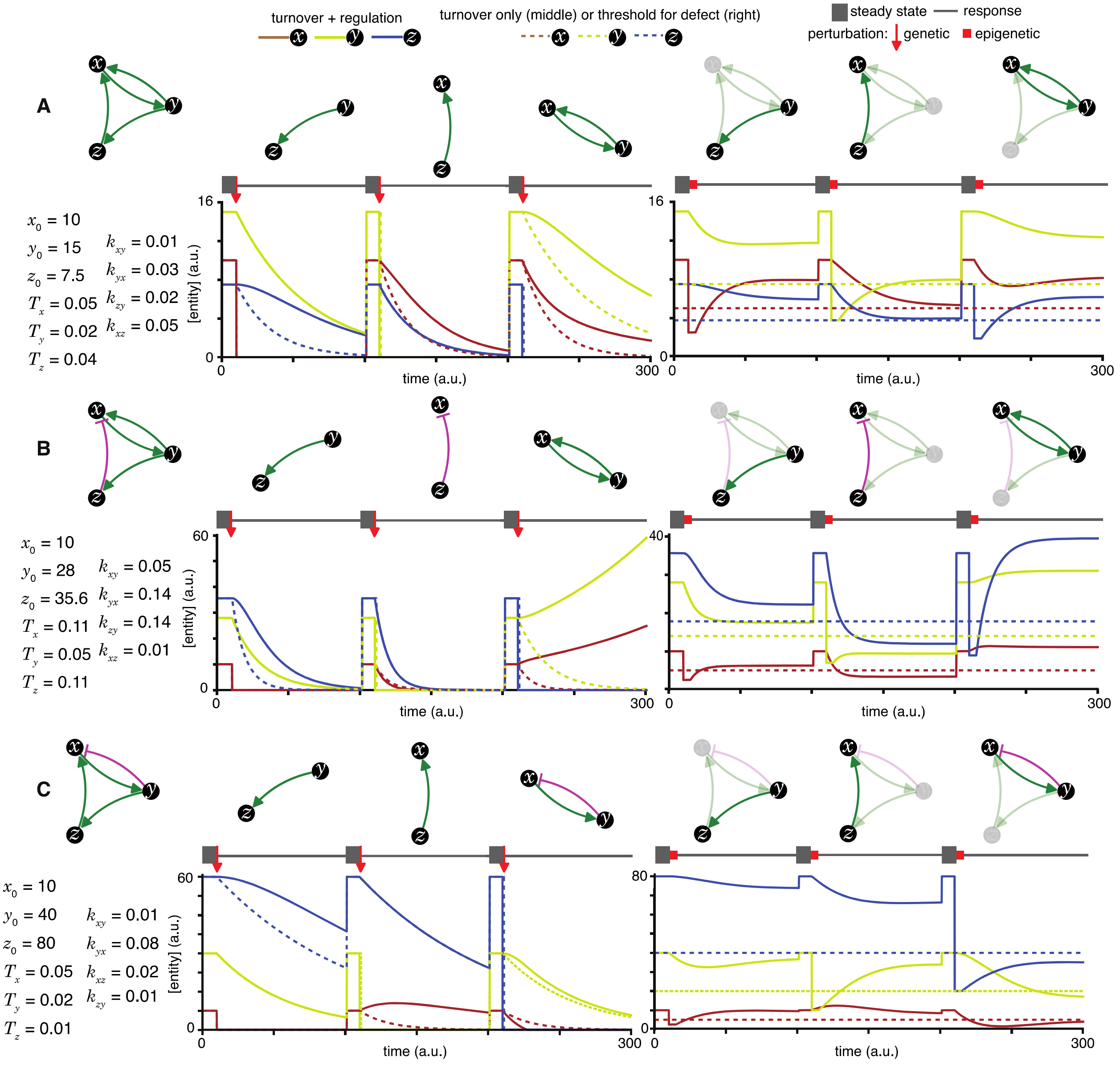


**Figure S6. Heritable regulatory architectures with two loops and a shared edge.** Architectures and their responses to perturbations are depicted as in Figure S4 (A, B, and C depict the heritable regulatory architectures H, I, and J, respectively).

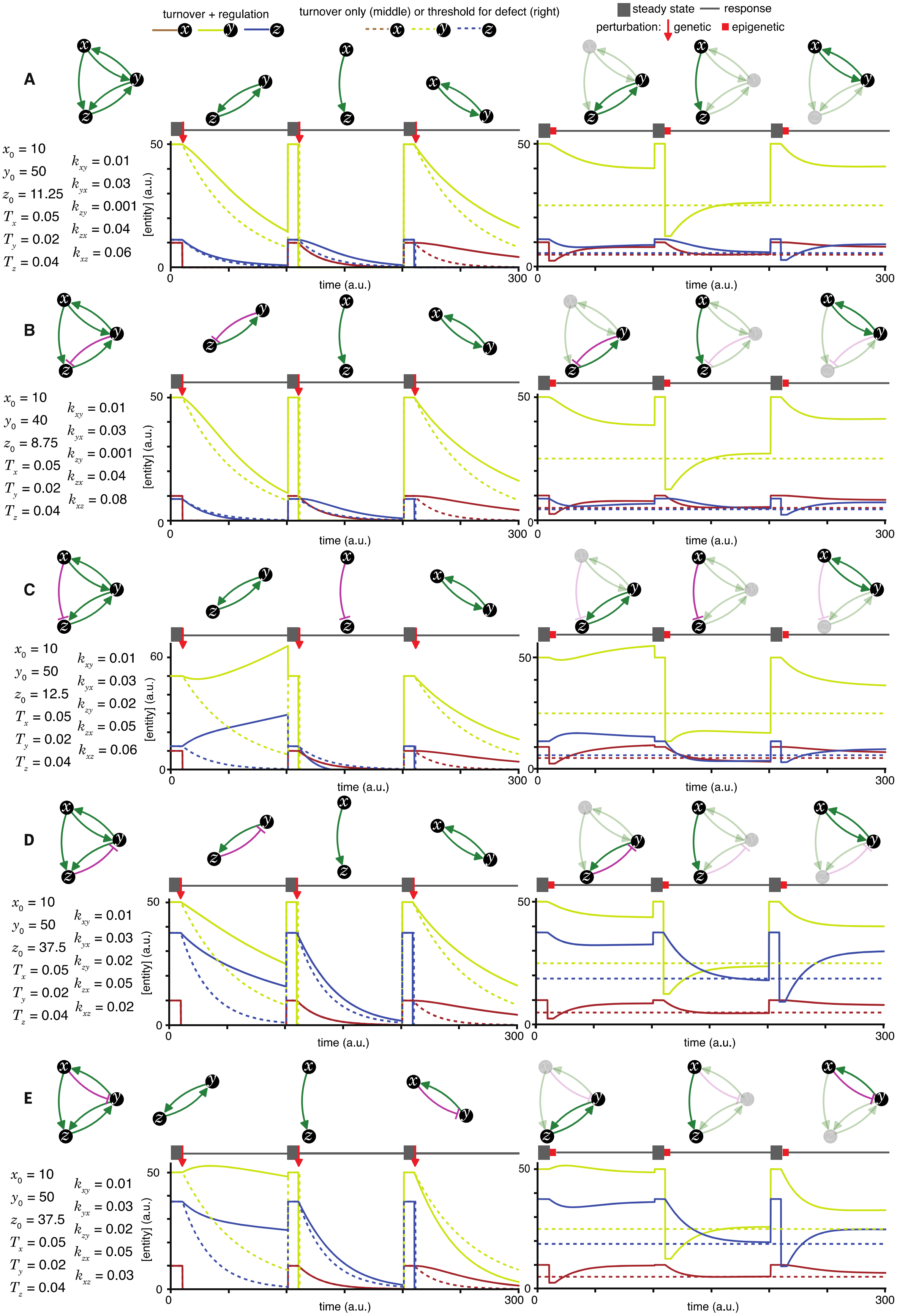

turnover + regulation
turnover only (middle) or threshold for defect (right)
steady state
response
perturbation: genetic
epigenetic
A
$x_0$ = 10
$y_0$ = 50
$z_0$ = 11.25
$T_x$ = 0.05
$T_y$ = 0.02
$T_z$ = 0.04
$k_{xy}$ = 0.01
$k_{yx}$ = 0.03
$k_{zy}$ = 0.001
$k_{zx}$ = 0.04
$k_{xz}$ = 0.06
B
$x_0$ = 10
$y_0$ = 40
$z_0$ = 8.75
$T_x$ = 0.05
$T_y$ = 0.02
$T_z$ = 0.04
$k_{xy}$ = 0.01
$k_{yx}$ = 0.03
$k_{zy}$ = 0.001
$k_{zx}$ = 0.04
$k_{xz}$ = 0.08
C
$x_0$ = 10
$y_0$ = 50
$z_0$ = 12.5
$T_x$ = 0.05
$T_y$ = 0.02
$T_z$ = 0.04
$k_{xy}$ = 0.01
$k_{yx}$ = 0.03
$k_{zy}$ = 0.02
$k_{zx}$ = 0.05
$k_{xz}$ = 0.06
D
$x_0$ = 10
$y_0$ = 50
$z_0$ = 37.5
$T_x$ = 0.05
$T_y$ = 0.02
$T_z$ = 0.04
$k_{xy}$ = 0.01
$k_{yx}$ = 0.03
$k_{zy}$ = 0.02
$k_{zx}$ = 0.05
$k_{xz}$ = 0.02
E
$x_0$ = 10
$y_0$ = 50
$z_0$ = 37.5
$T_x$ = 0.05
$T_y$ = 0.02
$T_z$ = 0.04
$k_{xy}$ = 0.01
$k_{yx}$ = 0.03
$k_{zy}$ = 0.02
$k_{zx}$ = 0.05
$k_{xz}$ = 0.03
[entity] (a.u.)
time (a.u.)

**Figure S7. Heritable regulatory architectures with two loops, a shared node, a connecting edge, and up to one negative regulatory interaction.** Architectures and their responses to perturbations are depicted as in Figure S4 (A, B, C, D, and E depict the heritable regulatory architectures K, L, M, N, and O, respectively).

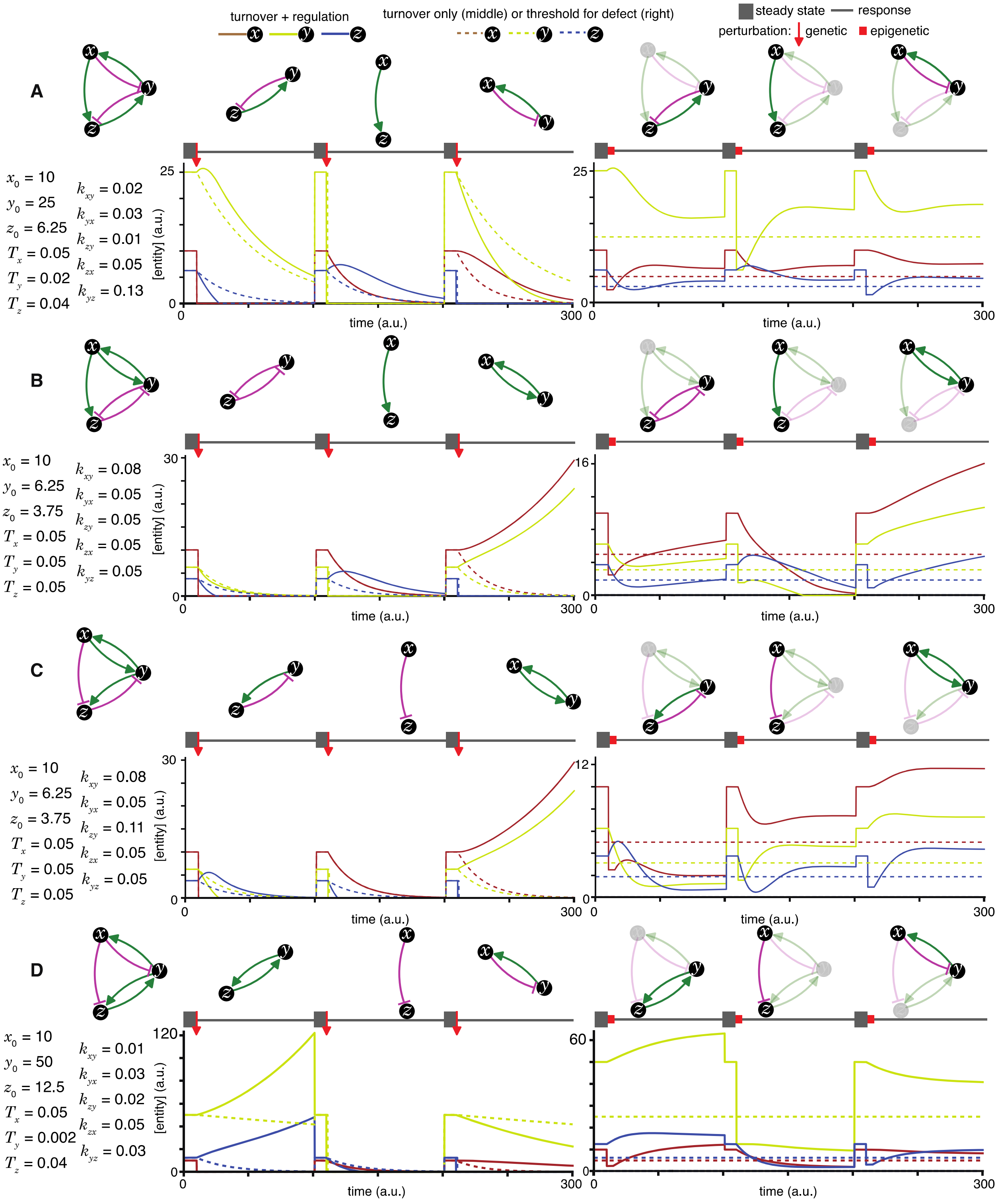


**Figure S8. Heritable regulatory architectures with two loops, a shared node, a connecting edge, and two negative regulatory interaction.** Architectures and their responses to perturbations are depicted as in Figure S4 (A, B, C, and D depict the heritable regulatory architectures P, Q, R, and S, respectively).

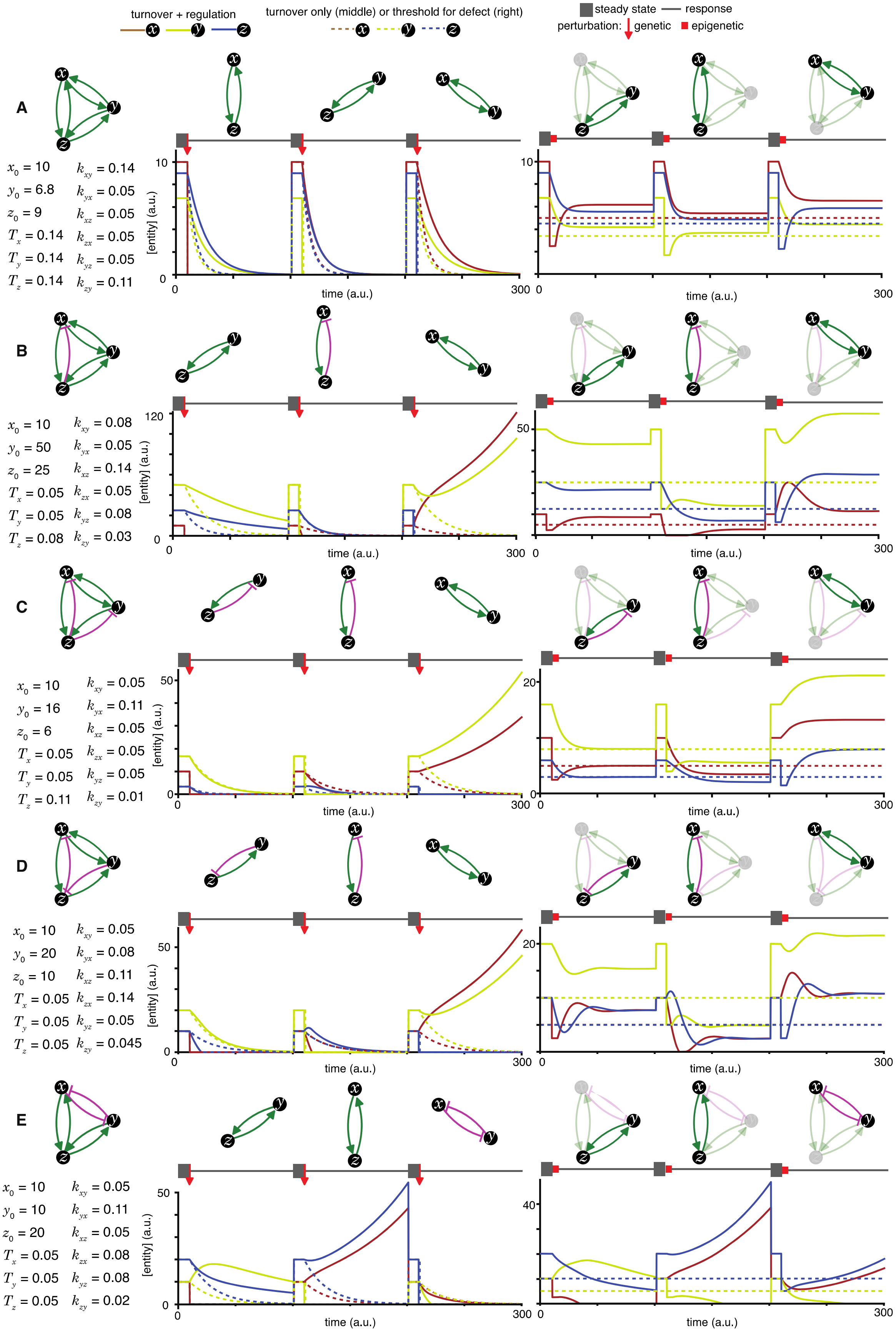

turnover + regulation
turnover only (middle) or threshold for defect (right)
steady state
response
perturbation:
genetic
epigenetic
A
$x_0$ = 10 $k_{xy}$ = 0.14
$y_0$ = 6.8 $k_{yx}$ = 0.05
$z_0$ = 9 $k_{xz}$ = 0.05
$T_x$ = 0.14 $k_{zx}$ = 0.05
$T_y$ = 0.14 $k_{yz}$ = 0.05
$T_z$ = 0.14 $k_{zy}$ = 0.11
B
$x_0$ = 10 $k_{xy}$ = 0.08
$y_0$ = 50 $k_{yx}$ = 0.05
$z_0$ = 25 $k_{xz}$ = 0.14
$T_x$ = 0.05 $k_{zx}$ = 0.05
$T_y$ = 0.05 $k_{yz}$ = 0.08
$T_z$ = 0.08 $k_{zy}$ = 0.03
C
$x_0$ = 10 $k_{xy}$ = 0.05
$y_0$ = 16 $k_{yx}$ = 0.11
$z_0$ = 6 $k_{xz}$ = 0.05
$T_x$ = 0.05 $k_{zx}$ = 0.05
$T_y$ = 0.05 $k_{yz}$ = 0.05
$T_z$ = 0.11 $k_{zy}$ = 0.01
D
$x_0$ = 10 $k_{xy}$ = 0.05
$y_0$ = 20 $k_{yx}$ = 0.08
$z_0$ = 10 $k_{xz}$ = 0.11
$T_x$ = 0.05 $k_{zx}$ = 0.14
$T_y$ = 0.05 $k_{yz}$ = 0.05
$T_z$ = 0.05 $k_{zy}$ = 0.045
E
$x_0$ = 10 $k_{xy}$ = 0.05
$y_0$ = 10 $k_{yx}$ = 0.11
$z_0$ = 20 $k_{xz}$ = 0.05
$T_x$ = 0.05 $k_{zx}$ = 0.08
$T_y$ = 0.05 $k_{yz}$ = 0.08
$T_z$ = 0.05 $k_{zy}$ = 0.02
[entity] (a.u.)
time (a.u.)
0
300

**Figure S9. Heritable regulatory architectures formed by complete graphs with up to two negative regulatory interactions.** Architectures and their responses to perturbations are depicted as in Figure S4 (A, B, C, D and E depict the heritable regulatory architectures T, U, V, W, and X respectively).

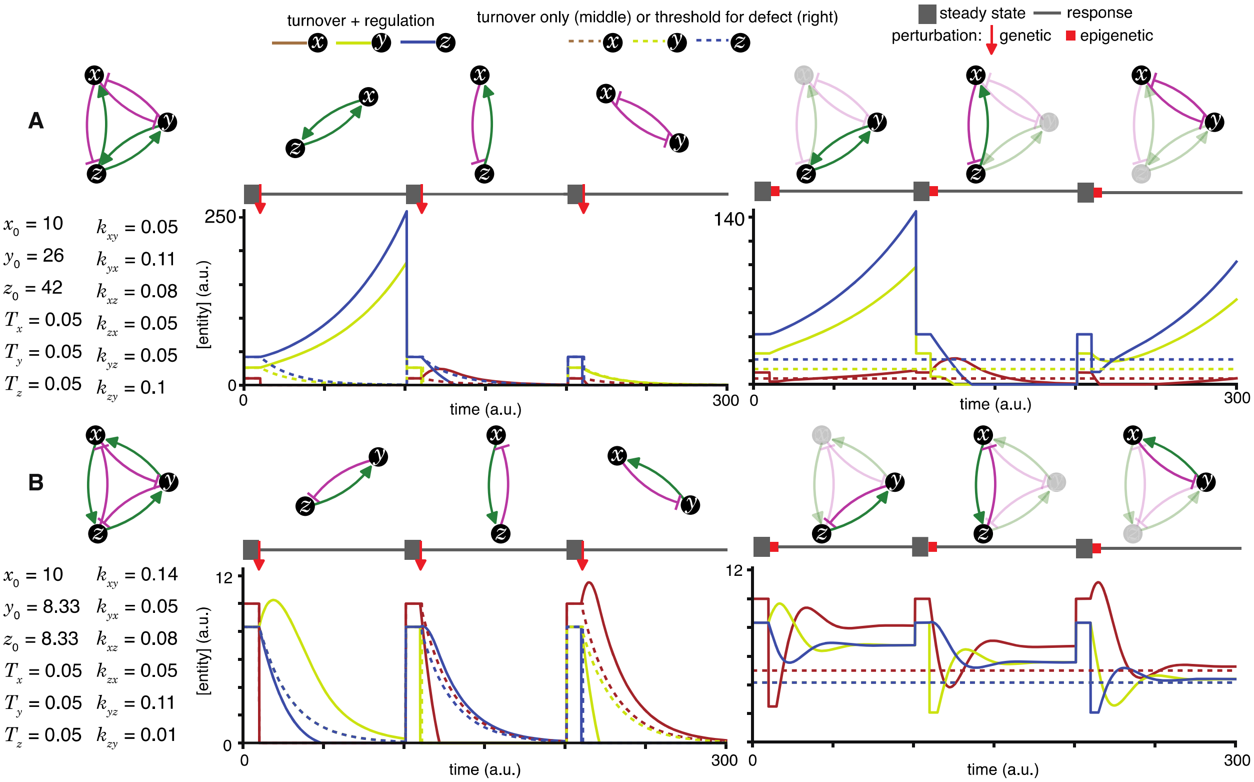


**Figure S10. Heritable regulatory architectures formed by complete graphs with three negative regulatory interactions.** Architectures and their responses to perturbations are depicted as in Figure S4 (A and B depict the heritable regulatory architectures Y and Z, respectively).

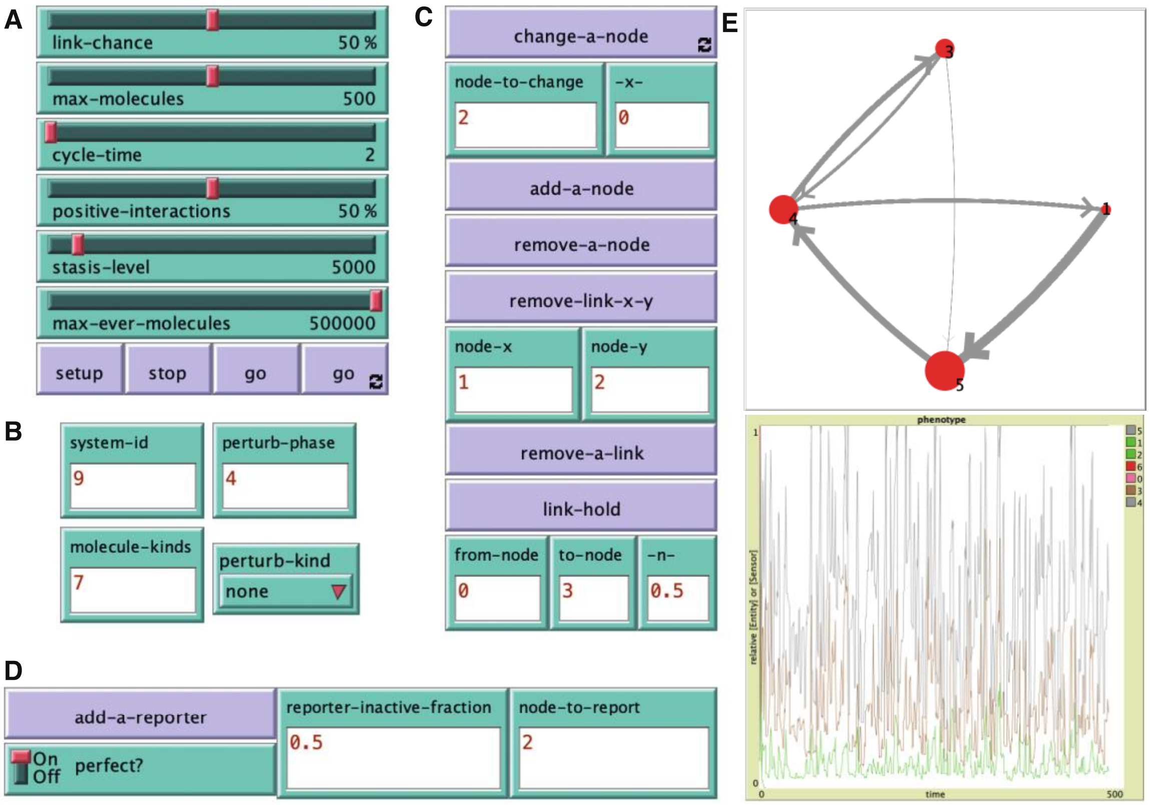


**Figure S11. Key features of the ESP system explorer.** A simulation of ESP systems was made using the agent-based modeling software NetLogo with controls for making a variety of changes. **(A)** Sliders and buttons for set up and simulation of an ESP system: link-chance, max-molecules, cycle-time, positive-interactions, stasis-level, max-ever-molecules, setup, stop, go, and go forever. **(B)** Parameters for specifying a particular system**:** system-id, molecule-kinds, perturb-phase, and perturb-kind. **(C)** Buttons and input for making changes to the system during the simulation**.** change-a-node, add-a-node, remove-a-node, remove-link-x-y (node-x, node-y), remove-a-link, and link-hold (from-node, to-node, -n-). **(D)** Buttons and input for adding a reporter of any node**.** add-a-reporter, perfect?, reporter-inactive-fraction, and node-to-report. **(E)** Representative output of changing regulatory architecture (*top*) and relative amounts of interactors representing 'phenotype' (*bottom*) over time. See Movie S1 for examples examining impact of changes and code (ESP_systems_single_system_explorer_v1.nlogo) for detailed information.

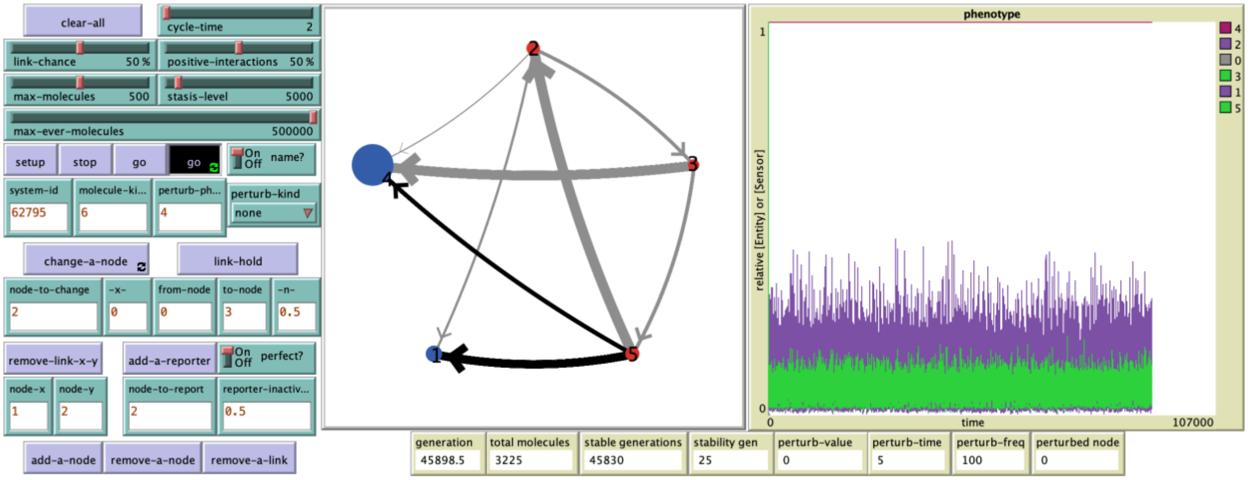

**Figure S12. Example of a system with long but finite stability.** This system (62795) begins with 6 entities/sensors, but after an early loss of one sensor, the remaining 5 are maintained as part of a HRA until 59,882.5 generations. See Fig. S10 and code (ESP_systems_single_system_explorer_v1.nlogo) for detailed information.

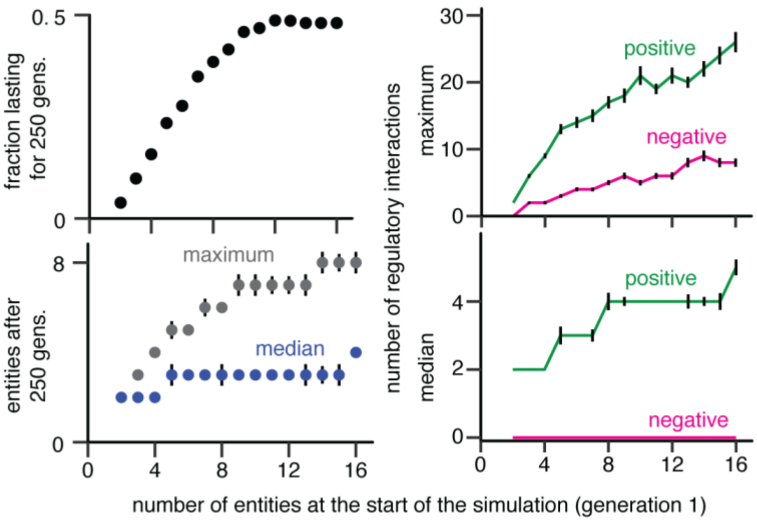


**Figure S13. Characteristics of randomly sampled HRAs simulated with partitioning of entities during each cell division or generation and periodic perturbations.** *Top left*, Fractions of ESP systems that persist with or without heritable epigenetic change for 250 generations when simulations were begun with different numbers of molecules. *Bottom left*, Maximum (grey) and median (blue) numbers of entities/sensors in ESP systems at the end of 250 generations when simulations were begun with different numbers of molecules. *Top right*, Maximal numbers of positive and negative regulatory interactions at the end of 250 generations when simulations were begun with different numbers of molecules. *Bottom right*, Median numbers of positive and negative regulatory interactions at the end of 250 generations when simulations were begun with different numbers of molecules.

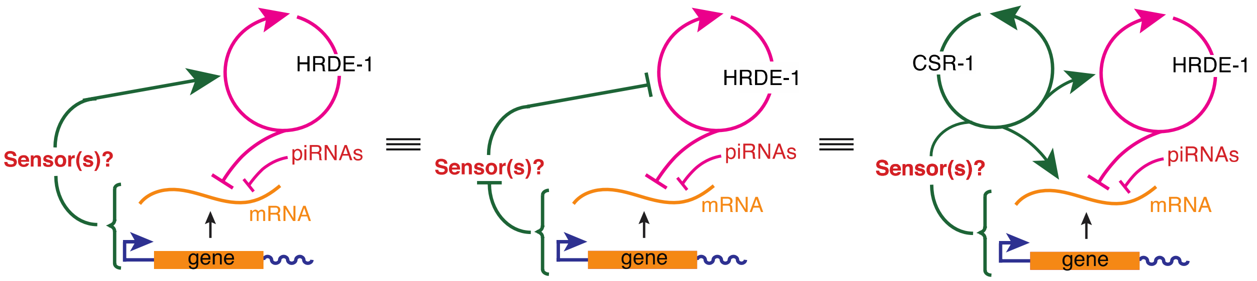


**Figure S14. Equivalent representations of a transgenerational feedback loop that can tune HRDE-1-dependent heritable RNA silencing.** *Left*, the architecture proposed in Fig. 6D, whereby a sensor(s) that promotes an HRDE-1-dependent positive feedback loop is reduced in response to changes in a gene or its gene products caused by the activity of the HRDE-1-dependent positive feedback loop, making it self-limiting. *Middle*, An architecture where the feedback from the gene to HRDE-1-dependent loop is via the inhibition of an inhibition instead of an activation. *Right*, An architecture as in *left*, but that includes additional CSR-1-dependent positive feedback loops that could amplify the transgenerational inhibition of the HRDE-1-dependent loop.

### SUPPLEMENTARY MOVIE LEGENDS

**Movie S1.** NetLogo run showing the single-system explorer with sample interactions with the simulation.

**Movie S2.** Example ESP system with system-id 46357 without any perturbation.

**Movie S3.** Example ESP system with system-id 46357 and with loss-of-function perturbations in phase 0.

**Movie S4.** Example ESP system with system-id 46357 and with loss-of-function perturbations in phase 1.

**Movie S5.** Example ESP system with system-id 46357 and with loss-of-function perturbations in phase 2.

**Movie S6.** Example ESP system with system-id 46357 and with loss-of-function perturbations in phase 3.

**Movie S7.** Example ESP system with system-id 46357 and with loss-of-function perturbations in phase 4.

**Movie S8.** Example ESP system with system-id 46357 and with gain-of-function perturbations in phase 0.

**Movie S9.** Example ESP system with system-id 46357 and with gain-of-function perturbations in phase 1.

**Movie S10.** Example ESP system with system-id 46357 and with gain-of-function perturbations in phase 2.

**Movie S11.** Example ESP system with system-id 46357 and with gain-of-function perturbations in phase 3.

**Movie S12.** Example ESP system with system-id 46357 and with gain-of-function perturbations in phase 4.

**Movie S13.** NetLogo run showing an ESP system with regulatory delays and developmental timing of cell divisions adapted from experimental results in *C. elegans*.
